# Diminishing effects of mechanical loading over time during rat Achilles tendon healing

**DOI:** 10.1101/2020.07.14.202077

**Authors:** Hanifeh Khayyeri, Malin Hammerman, Mikael J Turunen, Parmis Blomgran, Thomas Notermans, Manuel Guizar-Sicairos, Pernilla Eliasson, Per Aspenberg, Hanna Isaksson

## Abstract

Mechanical loading affects tendon healing and recovery. However, our understanding about how physical loading affects recovery of viscoelastic functions, collagen production and tissue organisation is limited. The objective of this study was to investigate how different magnitudes of loading affects biomechanical and collagen properties of healing Achilles tendons over time.

Achilles tendon from female Sprague Dawley rats were cut transversely and divided into two groups; normal loading (control) and reduced loading by Botox (unloading). The rats were sacrificed at 1, 2- and 4-weeks post-injury and mechanical testing (creep test and load to failure), small angle x-ray scattering (SAXS) and histological analysis were performed.

The effect of unloading was primarily seen at the early time points, with inferior mechanical and collagen properties (SAXS), and reduced histological maturation of the tissue in unloaded compared to loaded tendons. However, by 4 weeks no differences remained. SAXS and histology revealed heterogeneous tissue maturation with more mature tissue at the peripheral region compared to the center of the callus. Thus, mechanical loading advances Achilles tendon biomechanical and collagen properties earlier compared to unloaded tendons, and the spatial variation in tissue maturation and collagen organization across the callus suggests important regional (mechano-) biological activities that require more investigation.

## Introduction

Tendons are soft connective tissues responsible for load transmission with an energy-storing capacity that enables efficient locomotion. However, tendons recover poorly after injury, where the scar tissue that is formed has inferior biomechanical function. Tendons derive from mesenchymal stem cells and are mechanosensitive, meaning that tendons are affected by biophysical stimuli similar to other musculoskeletal tissues. However, how biophysical stimuli affects tendon healing is poorly understood, and this leaves many clinical treatments debated.

The Achilles tendon is the largest and the most commonly injured tendon in the body. Clinical reviews have shown that only 50% of patients with Achilles tendon ruptures regain pre-injury range of motion and load-bearing capacities. The dry weight of the tendon consists primarily of collagen (type I, 90% of dry weight) and its biomechanical function depends on composition and organisation. Collagen has typically a viscoelastic behaviour, which is characterised by specific time-dependent responses such as creep, stress-relaxation and hysteresis (Abrahams 1967; Cohen et al. 1976; Rigby et al. 1959). Each of these are believed to be connected to molecular interactions in collagen architecture (e.g. fibril alignment and sliding). However, the biomechanical regeneration during tendon repair is commonly evaluated with disruptive tests that mainly shed light on peak force (brittleness) and stiffness of the tissue. It does not evaluate the recovery of the essential time-dependent properties of tendons. Several studies have used dynamic loading to investigate tendon repair (Aubry et al. 2014; Eliasson et al. 2007; Freedman et al. 2018a; Freedman et al. 2018b). However, there are limited reports on how viscoelastic properties are restored during the time course of healing, and how loading affects this recovery process.

The collagen molecules are triple helixes that are staggered to form fibrils. At the fibril level the collagen appears with a periodicity, D-period, which is approximately 67 nm in mature healthy tendons. The D-period changes depending on the micromolecular environment, such as hydration (Turunen et al. 2017) or as a function of strain (Kukreti and Belkoff 2000). The fibrils are bundled into strands of collagen fibres, which are further packaged to make out the tissue. Tendon cells embedded in the collagen strands are mechanosensitive and respond to changes in their biophysical environment by altering their alignment (Buck 1980), signalling (Lavagnino et al. 2015; Wall et al. 2016) and synthetic activities (Chiquet et al. 2003; Killian et al. 2012). This modulates tissue composition and structure which in turn affects its biomechanical response to loading. It is however challenging to find the specific magnitude, frequency and timing of macroscopic loading that positively affects microscopic properties that in turn can be characterised as improved or accelerated repair and recovery.

Understanding how to use mechanical loading to regulate biological processes during tendon healing is invaluable for ultimately developing treatments that restore tendons to their post-injury state. Specifically, we need to better understand the recovery of the important viscoelastic properties of the tendon during repair, as the damping properties and the response of the tendon to dynamical loading is important.

The objective of this paper was to investigate how different magnitudes of daily loading affects the biomechanical and collagen properties of healing Achilles tendons over time. We hypothesised that reduced loading impedes collagen alignment and has a negative effect on the development of the viscoelastic properties over time.

## Methods

### Study design

Two separate but identical animal experiments were conducted; one for mechanical testing (N=10 in each group, 60 rats in total) and another for small-angle x-ray scattering (SAXS) and histological analysis (N=8 in each group, 48 rats in total) (Figure 1). In each animal experiment Sprague Dawley rats were divided into two groups: normal daily loading in the cage (full loading) or partial unloading where loading was reduced by Botox (unloading). The right Achilles tendon was transected and the rats were euthanized at 1, 2, or 4 weeks post-injury (Figure 1).

**Figure 1:**
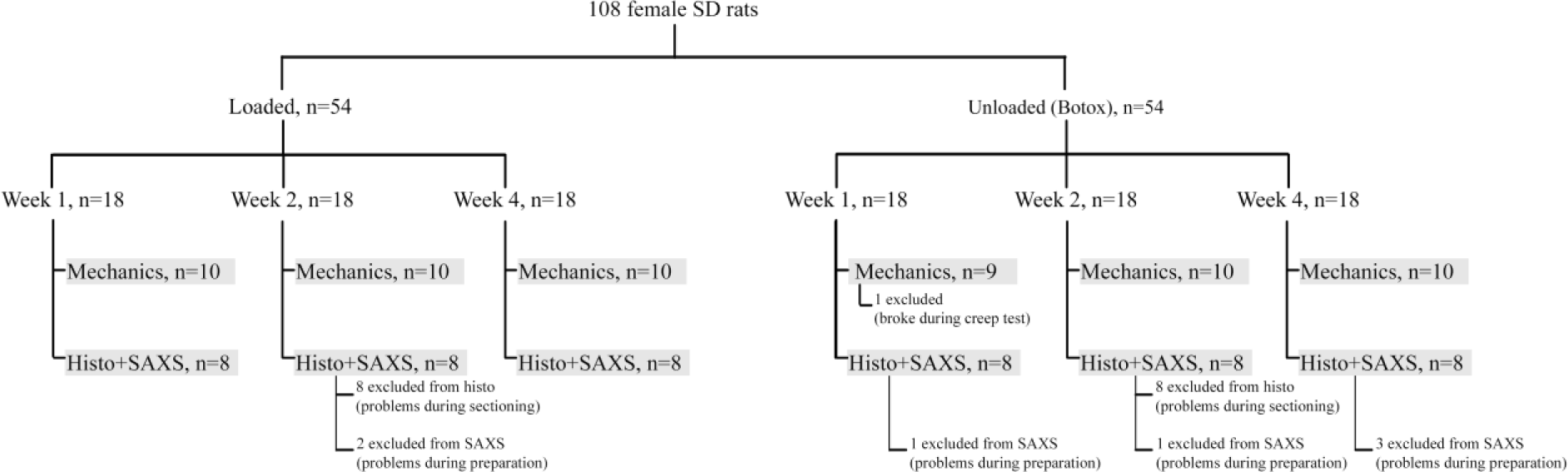
Experimental setup for the animal experiments indicating groups, times and number (n) of animals per group. Any exclusion of samples from the analyses is also indicated.

### Animal experiment

Female Sprague Dawley rats (aged 16 weeks mean weight 304 ±16 grams) were used for these experiments. All animals were randomly assigned to two different loading groups and the investigator was blinded during surgery and evaluation. In the unloaded group, the right gastrocnemius muscle (calf muscle) was injected with 1U Botulinum Toxin type A (Botox, Allergan, Irvine, CA) at three different sites (in total 3U/animal) 4 days before surgery. Visual inspection was done on all rats to confirm the effect of Botox injection. All rats underwent surgery: the right Achilles tendon were transversely transected in the middle of the tendon after removal of the plantaris tendon. The incision of the skin was sutured with two stiches and the tendons were left to heal spontaneously. Surgery and Botox injections were given under anaesthetics with Isoflurane gas (Forene, Abbot Scandinavia, Solna, Sweden). The animals received subcutaneous injections of antibiotics (Engemycin, 25 mg/kg oxytetracycline, Intervet, Boxmeer, the Netherlands) and analgesics (Temgesic, 0.045 mg/kg buprenorphine, Schering-Plough, Brussel, Belgium) preoperatively and analgesic (Temgesic) was given regularly until 48 hours after surgery. The transection day was considered as Day 0 in the experiments, and the animals were euthanized after 1, 2 or 4 weeks. All animals were allowed free cage activities from Day 0. All experiments were approved by the regional ethics committee for animal experiments in Linköping (Dnr 15-15) and adhered to the institutional guidelines for care and treatment of laboratory animals. The rats were housed two per cage and were given food and water *ad libitum*.

### Mechanical testing and characterisation

Tendons were harvested with the calcaneal bone and the gastrocnemius muscle. The callus size (medial-lateral and anterior-posterior diameters) was measured with a calliper before the mechanical test and the muscle was carefully scraped off avoiding the callus. Gap size was measured as the distance between the stumps using a calliper and placing the tendon in front of a strong light, which enabled clear visualization of the stumps inside the callus. The proximal tendon was put between sandpaper and clamped to the mechanical testing machine. The clamp on the proximal side was placed at the myotendinous junction that was visible by eye, and the distal tendon was clamped at the calcaneal bone. The angle between the tendon and the calcaneal bone corresponded to 30 degrees dorsiflexion.

The tendons were preconditioned by applying 10 cycles of loading between 1-2N at a rate of 0.1mm/s after which the tendons were allowed to rest approximately 4 hours in wet conditions. For all tendons, creep test was performed by applying 5N at a rate of 1mm/s, followed by holding the force constant for 300s. After unloading, the tendons rested for approximately 2-3 hours in wet environment. Subsequently, a second creep test was performed by applying 12N at a rate of 1mm/s, followed by holding the force constant for 300s. Finally, after a second period of resting, the tendons were loaded to failure at a rate of 1mm/s (Figure 2A). All tendons were tested within the same day of harvest. When resting between tests, the tendons were placed on a flat surface between wet gauze. The sandpaper was left in place to ensure that the clamps were placed on exactly the same place for the second creep test and load to failure.

**Figure 2:**
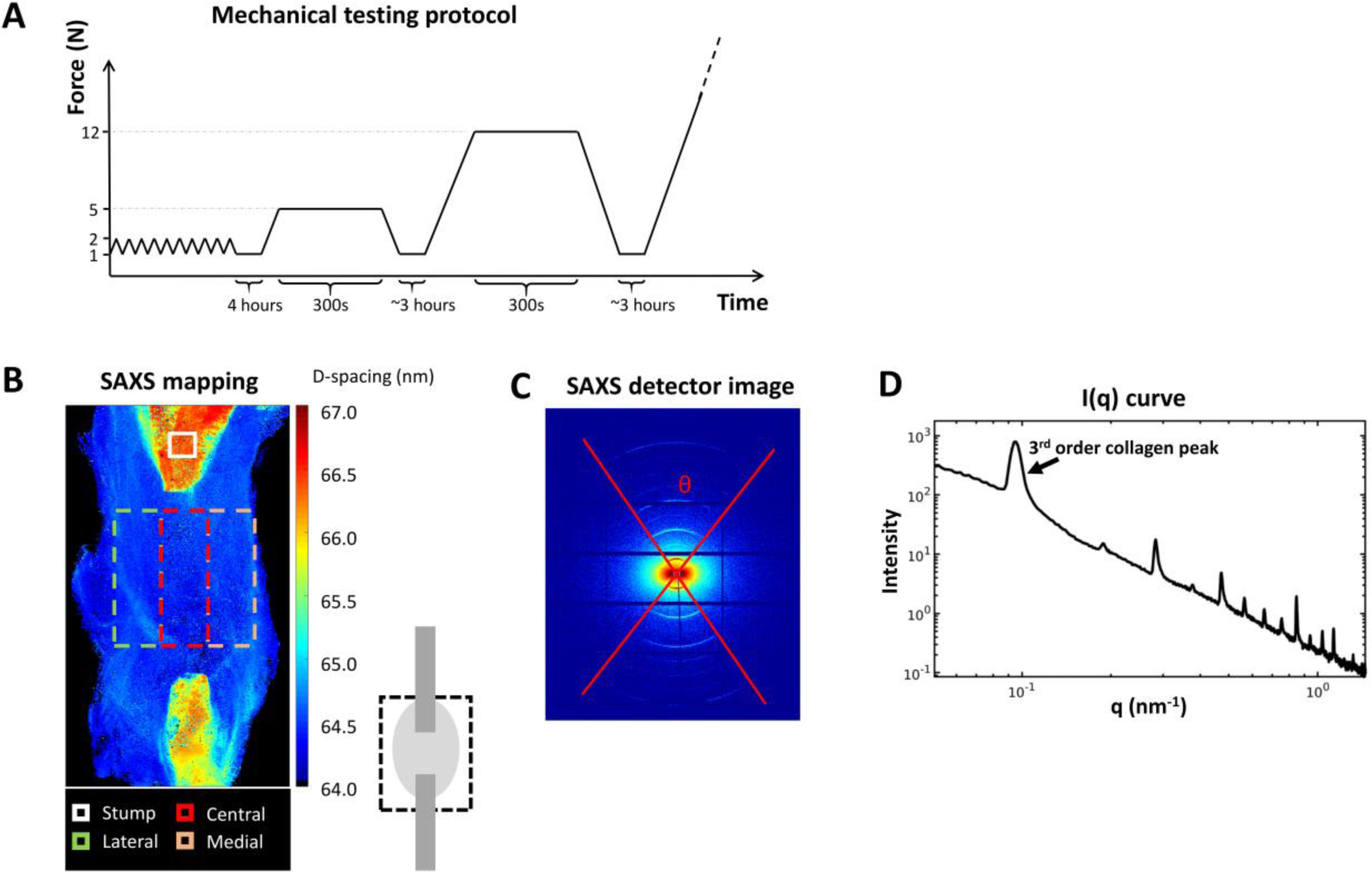
Analysis methods. A) Mechanical testing protocol included preconditioning, followed by two sets of creep tests before load to failure. B) Small Angle X-ray Scattering (SAXS) mapping over the tendon callus indicates the stumps and the regions of interest for analysis in the callus. C) SAXS detector image and D) integrated scattering intensity curve obtained from (C).

The following properties were analysed. Cross-sectional area was calculated by assuming an elliptical shape. Creep magnitude was measured as the displacement the tendon crept during the 300s holding phase. Creep ratio was calculated as the ratio between the creep magnitude and the gap size. The gap size was used instead of the clamping distance, as we predicted over 90% of the displacements to occur in the callus tissue initially. Tendon stiffness was calculated from the slope of the force-displacement curve at 80-90% of peak force in the creep tests (5N or 12N) and at 60-70% of the peak force in the ramp-to-failure test. The Young’s modulus was calculated as the slope from the stress-strain curve at similar locations. Upon failure, the peak force and peak stress were recorded from the force-displacement and stress-strain curves.

### Small-Angle X-ray Scattering (SAXS)

Tendons were harvested with calcaneus bone and gastrocnemius muscle. The muscle was carefully scraped off and the tendon was separated from the bone carefully. The tendon was pinned on silicone gels and covered with formaldehyde during transport and until analysis. The time between harvest and analysis was less than 14 days. To keep the samples moist during SAXS measurements, the tendons were placed in small handmade Kapton pockets that were filled with formaldehyde. The Kapton pockets were taped to the sample holder for the measurements. All SAXS measurements were made on the whole tendons (i.e. not sectioned).

The cSAXS beamline at the Swiss Light Source (SLS), Paul Scherrer Institute (PSI), Villigen, Switzerland was used with a Pilatus 2M detector (16) recording the 2D scattering patterns of the tendons in a continuous line-scan mode (raster-scan with 30 x 30 µm^2^ spot size and step size of 30 µm). Exposure time of 50 ms and X-ray wavelength of 1.0 Å was used. The sample-detector distance was 7108 mm giving a *q* range of ∼0.02-1.45 nm^-1^. Silver-Behenate (AgBH) powder standard was used to determine beam center and sample-detector distance. A rectangular region in the mid-region of the callus was selected, using a camera with a calibrated distance to the X-ray beam, for measurement and analysis (Figure 2B). This region was divided into five areas horizontally: medial, central, lateral, peripheral callus and full tendon, the latter consisting of the first three regions (Figure 2B). Additionally, a background line-scan next to each sample on the respective the Kapton pocket was recorded. The data were background corrected by removing the background scattering from the tendon scattering.

The scattering data contains information about the collagen structure and orientation. In this study, five collagen parameters were analysed according to previous protocols (Khayyeri et al. 2017). The anisotropy of collagen fibres (θ, in degrees) was determined as the width of the collagen rings in the scattering image (Figure 2C), from the full width of the tenth maximum. Peak location was measured as the radii of the 3^rd^ concentric arc in the scattering image where D-spacing (periodicity) was inversely proportional to peak location. By azimuthally integrating the detector images over theta (θ), the intensity, I(q), of the scattering images was obtained containing structural information (Figure 2D). From the intensity curve (I(q)), the third order collagen peak was analysed by fitting a Gaussian curve. Peak intensity (fibre alignment), full width at half maximum (FWHM, fibre delamination), and peak area (interfibrillar ordering) were calculated from the Gaussian fit.

Profiles of the variation of the different parameters across the tendons were calculated by averaging the parameter maps within the ROI vertically (Figure 2B). For comparison between the samples and further group-wise averaging, the lengths of the profiles were normalized by interpolating all profiles to the same length. All analyses were done with in-house written scripts in Matlab (R2016b) (Turunen et al. 2017).

### Histology

After SAXS measurements, the samples were prepared for histology following a standard protocol with ethanol dehydration and paraffin embedding. Embedded tendons were sectioned longitudinally to 3µm thick slices and stained with Hematoxylin & Eosin, Picrosirius red for collagen, and Alcian blue (pH 2,5) for glycosaminoglycans (GAGs). Sections were taken where both the tendon stumps and the mid callus region was visible. All sections with the different staining methods were observed using a light microscope (Zeiss Axio). The data presented is descriptive.

A semi-quantitative analysis for tissue maturation was performed on sections stained with Hematoxylin & Eosin. Three images were captured from each section in the middle of the callus in between the stumps (a medial, central and lateral image with magnification 20X). A blinded investigator graded all images between 1 and 4 for cell number, nuclear shape, collagen alignment and collagen stainability where a low score represented a more mature tendon (Figure 3). Scoring description: Cell number: 1 = relatively few cells and 4 = high number of cells. Nuclear shape: 1 = most of the cells are spindle shaped, 2 = many of the cells are spindle shaped, 3 = some of the cells are spindle shaped and 4 = very few cells are spindle shaped. Collagen alignment: 1 = almost all collagen in the picture is aligned in a parallel manner, 2 = more than 50% of the collagen is aligned, 3 = less than 50% of the collagen is aligned and 4 = difficult to see a direction of the collagen alignment. Collagen stainability: 1 = intense staining (dark pink) and 4 = weak staining (light pink).

**Figure 3:**
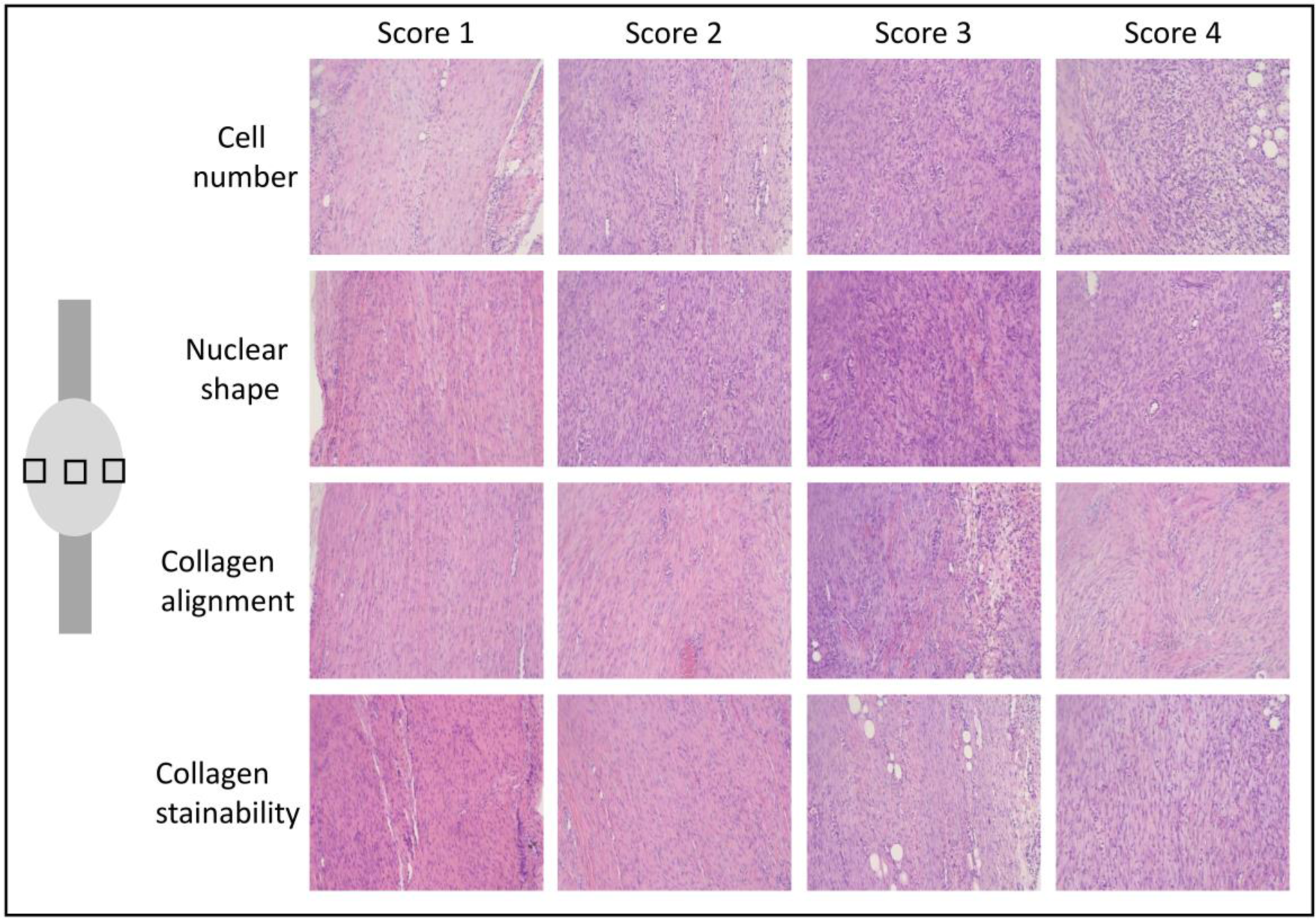
Histological evaluation. Pictures from the semi-quantitative analysis for tissue maturation of healing tendons at 1 and 4 weeks of healing. Three images were captured from each specimen in the middle of the callus in between the stumps as shown in the picture at the left side. The specimens were stained with hematoxylin & eosin and the picture was taken with a magnification of 20. A blinded investigator graded all pictures between 1 and 4 for cell number, nuclear shape, collagen alignment and collagen stainability and a representative picture from each score is shown. A low score represents a more mature tendon.

### Statistics

Statistical comparison was performed with a two-way ANOVA to assess the effect of treatment (loading and unloading) and healing time (1, 2, 4 weeks). Significant relationships (p ≤ 0.05) were further analysed with post-hoc Student’s t-test correcting for multiple groups when needed. For histology, also repeated measures with respect to location was added to the model. All data was analysed in IBM SPSS version 23.

## Results

### Mechanical testing

Overall, there were geometrical differences between the loaded and unloaded group. The cross-sectional area in the loaded group was largest at 1 week post-injury and showed a decreasing trend over-time. The cross-sectional area was higher in the loaded group at 1 week compared to the unloaded group, but the difference decreased over-time to be similar at 2 and 4 weeks post-injury (Figure 4A). The gap size in the loaded and unloaded group remained similar over time, but was larger in the loaded group compared to the unloaded group at 1 and 2 weeks post-injury (Figure 4B).

**Figure 4:**
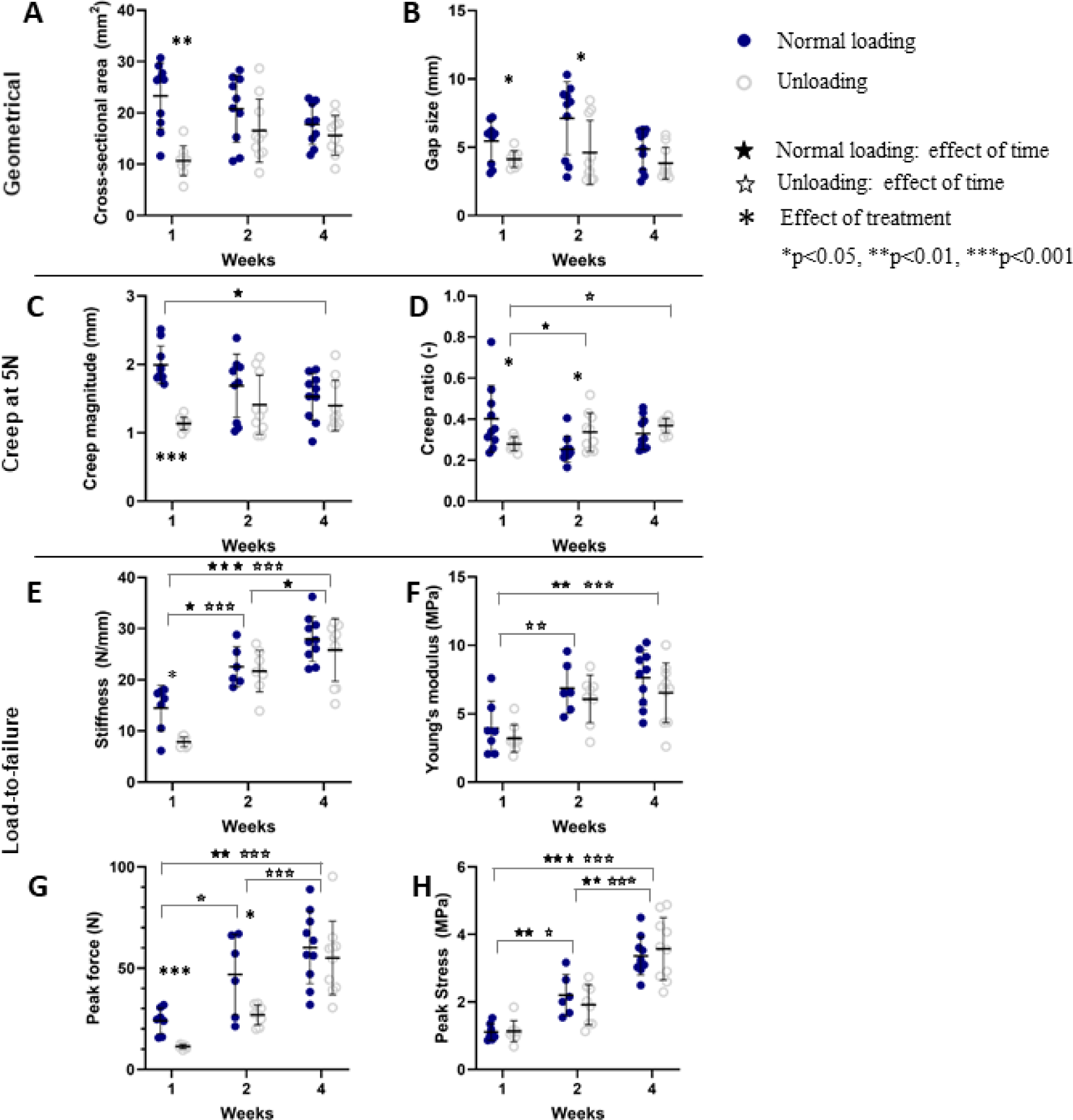
Geometrical and mechanical properties in loaded and unloaded healing tendons measured after 1, 2 and 4 weeks of healing. The callus geometry is described by cross-sectional area and gap size. Creep magnitude and ratio from the 5N creep test are shown. From the tension-to-failure test, stiffness, Young’s modulus, peak force and peak stress are displayed. Individual data points, as well as mean and standard deviation are shown in the graphs.

Creep magnitude and creep ratio was higher at 1 week post-injury in the loaded group as they exhibited more pronounced creep behaviour (Figure 4 C-D; see Suppl. table 1). The creep magnitude reduced over time in the loaded group while it remained similar in the unloaded group, such that both groups showed a similar creep behaviour at 4 weeks post-injury (Figure 4C). The creep ratio increased in the unloaded group over time, whereas it decreased over time in the loaded group (Figure 4D).

**Table 1:** Semi-quantitative analysis in healing tendons at 1 and 4 weeks.

|  | Analysis | Treatment | Lateral |  | Centre |  | Medial |  | Σ of all pictures |
| --- | --- | --- | --- | --- | --- | --- | --- | --- | --- |
| 1 week | Cell number | Full loading | 2 | (2-3) + | 3 | (2-4) | 3 | (2-4) | 8 (7-11) |
|  |  | **Botox** | **2.5** | **(2-4)** | **3** | **(2-4)** | **3.5** | **(3-4)** | **10 (8-11)** |
|  | Nuclear shape | Full loading | 3 | (3-3) | 3 | (2-3) | 3 | (2-3) | 9 (7-9) |
|  |  | **Botox** | **3** | **(2-4)** | **3.5** | **(2-4)** | **3** | **(3-4)** | **9 (8-11)** |
|  | Collagen alignment | Full loading | 2 | (2-3) | 3 | (1-3) | 2 | (2-3) | 7 (5-9) * |
|  |  | **Botox** | **2.5** | **(2-3)** | **4** | **(2-4)** | **3** | **(2-4)** | **9 (7-10)** |
|  | Collagen stainability | Full loading | 2 | (2-3) + ‡ | 4 | (2-4) | 3 | (3-4) | 7 (5-9) |
|  |  | **Botox** | **3** | **(2-4)** | **3** | **(3-4)** | **3** | **(2-4)** | **9 (7-10)** |
|  | Maturation score | Full loading | 10 | (9-11) + ‡ | 11 | (10-13) | 12 | (10-13) | 32 (29-37) * |
|  |  | **Botox** | **10.5** | **(9-14) +** | **14** | **(10-16)** | **12.5** | **(11-15)** | **36.5 (31-41)** |
| 4 weeks | Cell number | Full loading | 2 | (1-3) | 2 | (1-2) | 2 | (1-3) | 6 (3-8) * |
|  |  | **Botox** | **3** | **(1-4)** | **2.5** | **(1-4)** | **2.5** | **(1-3)** | **7.5 (6-10)** |
|  | Nuclear shape | Full loading | 2 | (1-4) | 2.5 | (2-4) | 2.5 | (1-3) | 7 (5-10) |
|  |  | **Botox** | **3** | **(1-4)** | **4** | **(2-4)** | **3** | **(2-3)** | **9.5 (6-11)** |
|  | Collagen alignment | Full loading | 2 | (1-4) | 2.5 | (1-4) | 2.5 | (1-4) | 7.5 (4-11) |
|  |  | **Botox** | **2.5** | **(1-4) +** | **4** | **(1-4)** | **3** | **(2-3)** | **9 (5-10)** |
|  | Collagen stainability | Full loading | 2.5 | (1-3) | 2.5 | (1-3) | 3 | (1-4) | 7.5 (3-10) |
|  |  | **Botox** | **2** | **(1-3)** | **2.5** | **(1-4)** | **2** | **(1-3)** | **6 (4-9)** |
|  | Maturation score | Full loading | 8 | (6-14) | 9.5 | (6-12) | 9 | (7-11) | 28 (19-35) # |
|  |  | **Botox** | **10.5** | **(6-15)** | **13** | **(7-15)** | **10** | **(7-11)** | **32.5 (24-35) #** |

The numbers are shown as median and range. A low score represents a more mature tendon. A lower score means less cells, less rounded shape, better collagen orientation and more stained collagen.
\* significant difference between full loading (normal) and unloading by botox.

Stiffness and Young’s modulus were lower in the loaded tendons compared to the unloaded ones during the 5N creep load at 1 week post-injury (see Suppl. table 1) but in contrast, the stiffness measured during load to failure at 1 week post-injury was higher in the loaded tendons (Figure 4E). In the load to failure, stiffness and Young’s modulus increased significantly over time in both treatment groups (Figure 4E-F). The Young’s modulus was similar for the loaded and unloaded tendons throughout all time points. Peak force (ultimate strength) and peak stress (ultimate stress) increased substantially over time in both groups (Figure 4 G-H). In terms of ultimate strength, the loaded tendons displayed a higher peak force than the unloaded tendons at 1 and 2 weeks (Figure 4 G).

### SAXS

Examples of spatial development over time in loaded and unloaded tendons are visible in Figure 5. Analysis of D-spacing showed that the unloaded group exhibited shorter D-spacing compared to the loaded group at 1 and 2 weeks post-injury (Figure 6A). D-spacing increased over time throughout the tendon and in both groups. After 4 weeks of healing, there was no difference between the loaded and unloaded tendons. The D-spacing was then approximately 65 nm (Figure 6A, Supplementary Figure 2). This is an increase with about 0.5 nm from 1 week post-injury but still 2 nm below the average values of D-spacing in healthy (uninjured) collagen fibrils in rat Achilles tendon (Khayyeri et al. 2017).

**Figure 5:**
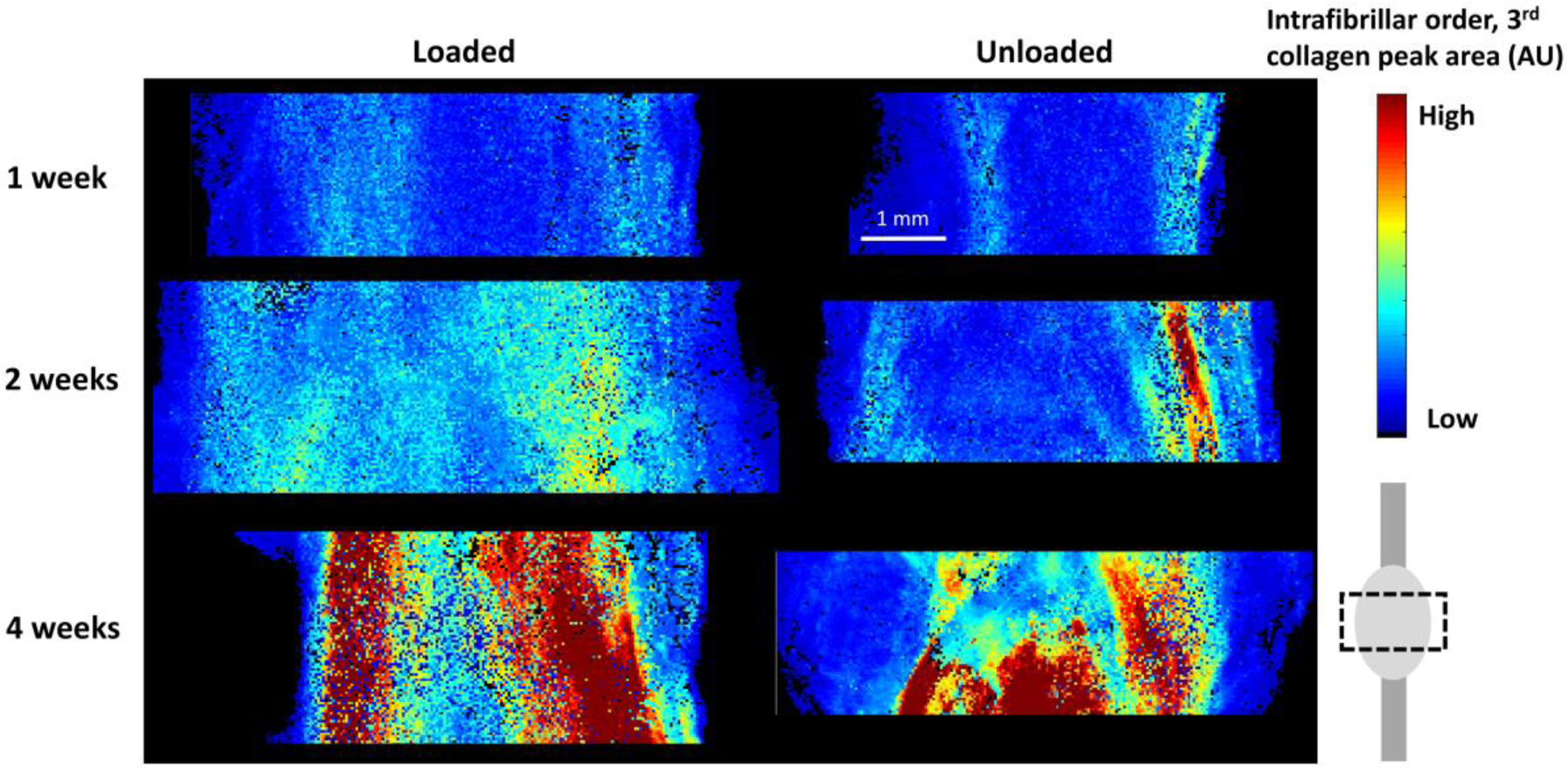
SAXS mapping of the intrafibrillar order parameter (peak area, AU) in the callus region in representative loaded and unloaded samples at 1, 2 and 4 weeks of healing.

**Figure 6:**
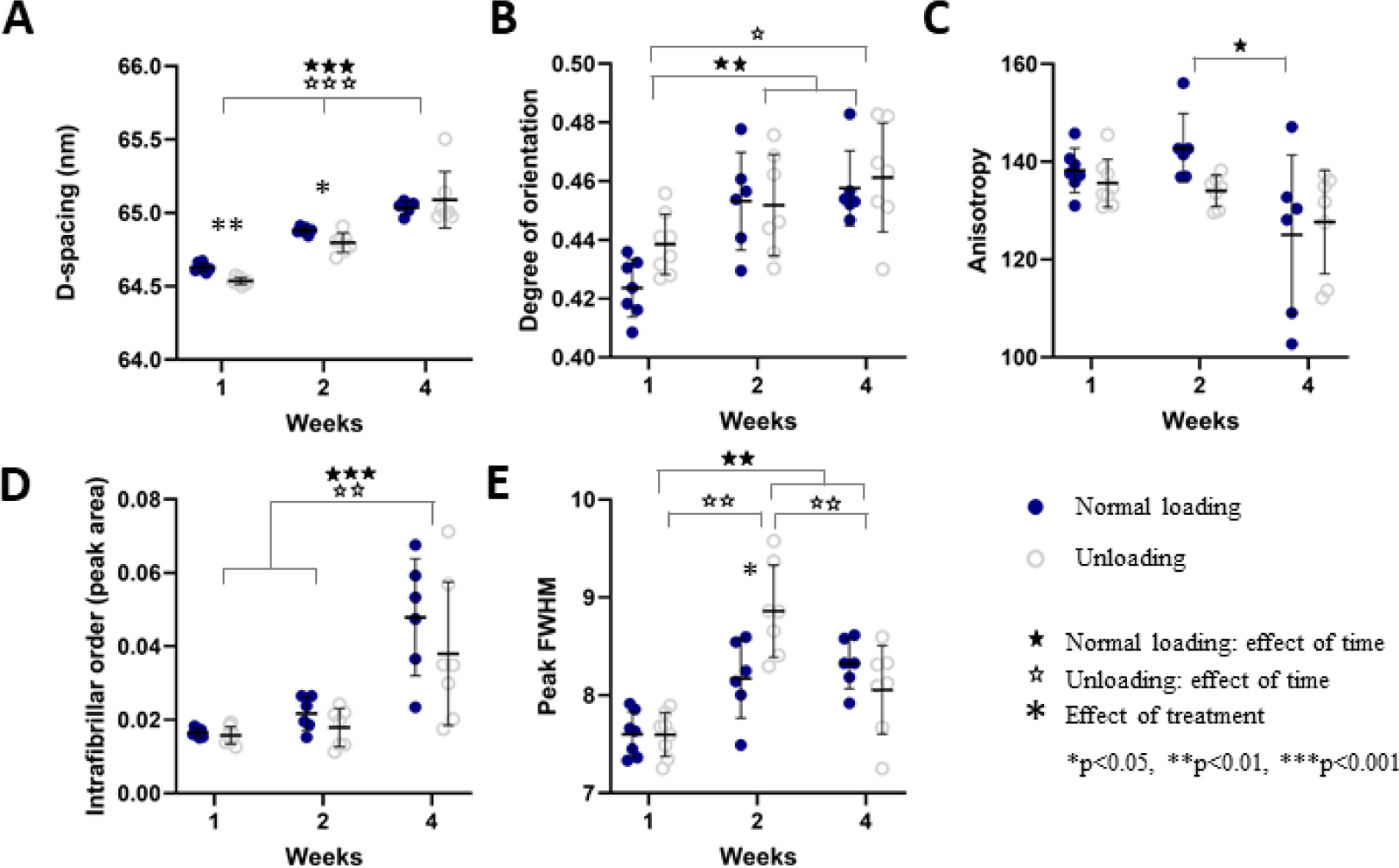
Quantitative SAXS data from the callus region for loaded and unloaded samples after 1, 2 and 4 weeks of healing, displaying A) D-spacing (nm), B) Degree of orientation (AU), C) anisotropy (degrees), D) intrafibrillar order (AU) and E) peak FWHM (nm). Individual data points, as well as mean and standard deviation are shown in the graphs.

Fibril alignment, measured from peak intensity, revealed no difference in degree of orientation between the loaded and unloaded group (Figure 6B), but the alignment increased over time in both groups.

There was no remarkable difference in degree of anisotropy (measured as the dispersion angle in the scattering image) between the two groups at any of the time points (Figure 6C). From the profiles, it was observed that anisotropy seemed to be lower in the medial and lateral side of the tendon callus compared to the centre (Figure 7C). This is more clearly reflected in the intrafibrillar order where organisation visibly increases over the course of healing (Figure 6D, 7D). The effect of loading was only apparent at the centre of the tendon, showing higher intrafibrillar organisation in the loaded group compared to the unloaded group after 1 and 2 weeks of healing (Supplementary table 2).

**Figure 7:**
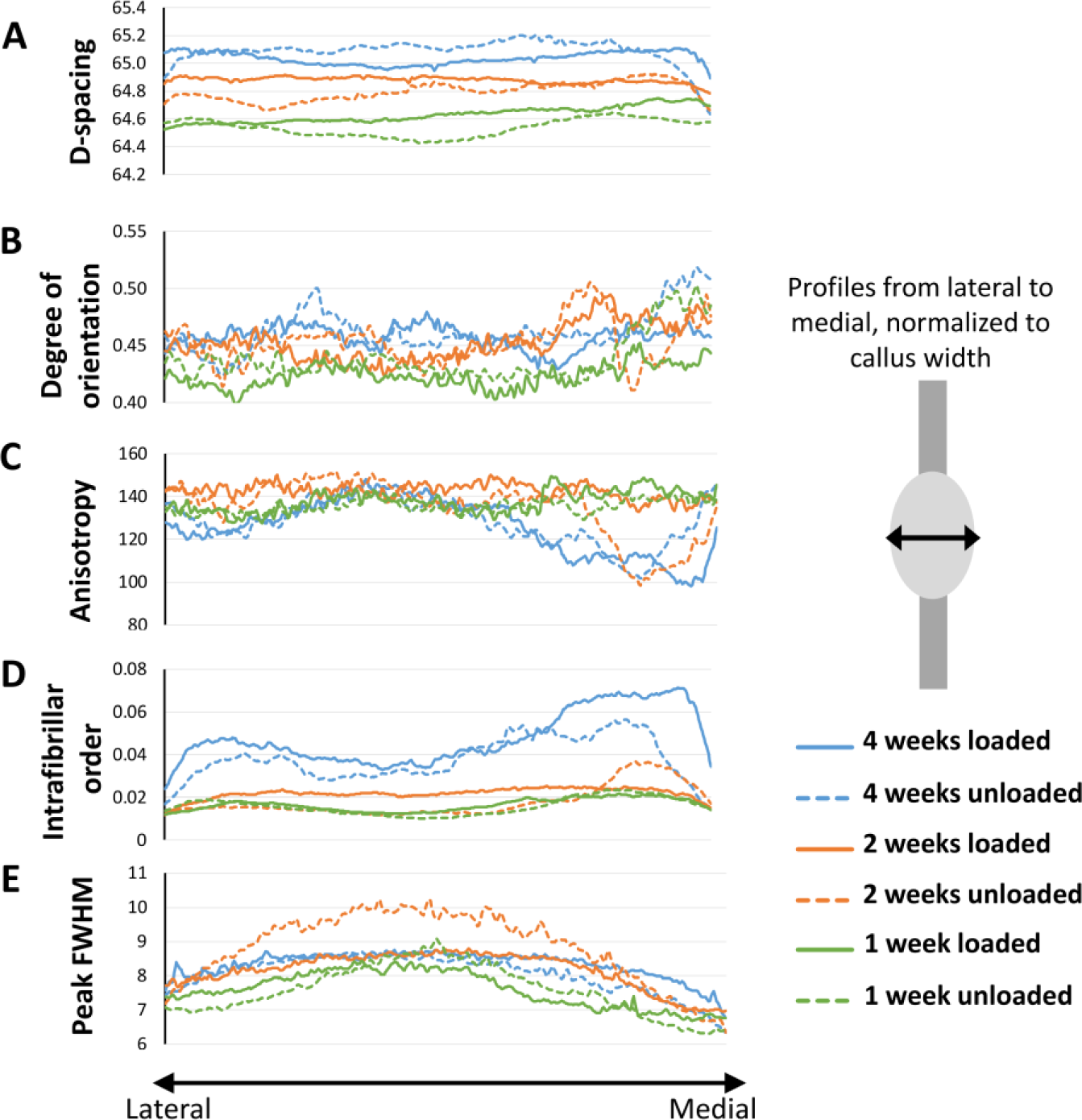
Spatial profiles of the SAXS data from the lateral (left) to the medial (right) side of the callus for loaded and unloaded samples after 1, 2 and 4 weeks of healing, displaying A) D-spacing (nm), B) Degree of orientation (AU), C) anisotropy (degrees), D) intrafibrillar order (AU) and E) peak FWHM (nm). Average profiles of all samples per group are shown.

Full width at half maximum (FWHM) measures collagen fibril adhesion and packing to some degree, where higher FWHM means less fibril adhesion and looser packing of the collagen. There was no conclusive difference between the treatment groups, but an increase in FWHM was observed 2-weeks post-injury in the unloaded group, which was reduced again by week 4. FWHM increased over time for both loaded and unloaded groups. (Figure 6E). The profiles showed that FWHM was higher in the centre of the callus compared to the medial and lateral sides. This pattern was clear at 1 and 2 weeks post-injury in both groups (Figure 7E). All quantitative analysis in separate regions are available in Supplementary table 2.

### Histology

The dichotomy caused by gap size seen in the mechanical testing experiment was also visible in the histology, i.e. the gap size was small in the unloaded group and larger in the loaded group. Therefore, 3 images of newly formed tissue in different regions (medial, central and lateral) were captured for a semi-quantitative analysis.

Overall, the semi-quantitative analysis showed that loaded tendons were significantly more mature at 1 week compared to the unloaded tendons (lower score reflects a more mature tissue), while there was no significant difference at 4 weeks of healing (Table 1). The loaded tendons had a lower score in nearly all parameters (cell number, nuclear shape, collagen alignment and collagen stainability) compared to the unloaded tendons at 1 week, but only the difference in collagen alignment was significant. Time improved the score more for the unloaded tendons, hence there was no significant difference in the total maturation score between the loaded and unloaded group at 4 weeks. The cell number seemed to be reduced with time in both groups (Table 1), while nuclear shape was only improved in loaded tendons and collagen stainability was only improved in the unloaded tendons (Table 1). The callus tissue appeared to mature with a regional variation. The three regions in the loaded tendons was significant different regarding the maturation score at 1 week where the lateral side appeared to contain the most mature tissue and then the maturation reduced continuously to the other side (Table 1). However, this discrepancy at the different regions in loaded tendons was fully diminished after 4 weeks. The unloaded tendons seem to heal in a different manner having more mature tissue at the sides and less mature in the centre, especially in terms of collagen alignment and nuclear shape. These differences were not statistically significant although this trend was seen both at 1 and at 4 weeks of healing.

The two other staining’s (Picrosirius red and Alcian blue) were thoroughly observed in a qualitative and descriptive manner. Adipocyte were seen within the callus tissue in both groups at 1 week, but it was more common in the loaded tendons (Figure 8 A vs C). However, the unloaded tendons had more adipose cells at the sides of the callus (Figure 8 C). At 4 weeks, there was less adipocytes in the callus in both groups (Figure 8 E and G). Although, most of the unloaded tendons exhibited a layer of adipocytes on one side of the callus (Figure 8 G), i.e. the side that had less collagen alignment, whereas the loaded tendons had a low adipocyte number on both sides of the callus. Bleeding in the callus, seen as extravasated erythrocytes, was found to a varied degree in several tendons from both groups at 1 week of healing but this could not be seen at 4 weeks. The tendon stumps at 4 weeks were less distinct in both groups. The Alcian blue staining revealed more proteoglycans only in the proximity of the stumps in the unloaded tendons at 1 week (Figure 8 K) while the loaded tendons had more regions throughout the callus with more proteoglycans (Figure 8 I). After 4 weeks of healing, the amount of proteoglycans was reduced in the callus in the unloaded tendons in most of the samples, but area around the distal stump contained more proteoglycans as well as chondrocytes in almost all samples (Figure 8 O). The loaded group again showed a heterogeneous distribution of “islands” with proteoglycans throughout the whole callus tissue while most chondrocytes were localised close to the tendon stumps (Figure 8 M).

**Figure 8:**
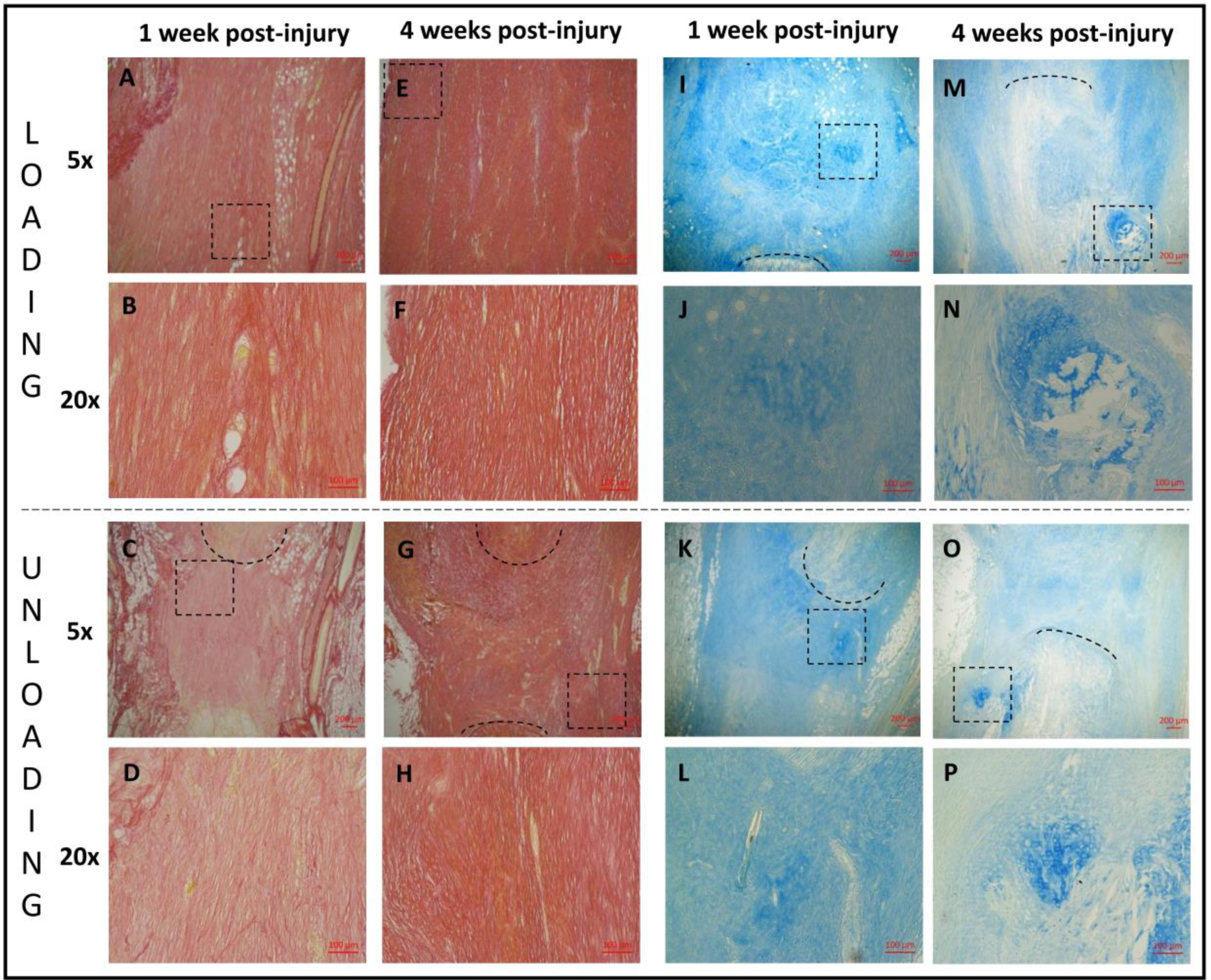
Histology. Healing tendons from 1 (A, B, C, D, I, J, K, L) and 4 weeks (E, F, G, H, M, N, O, P) post-injury stained with picrosirius red (A, B, C, D, E, F, G, H) or alcain blue (I, J, K, L, M, N, O, P). Healing tendons exposed to loading (A, E, I, M (magnification 5) and B, F, J, N (magnification 20)) can be compared to healing tendons exposed to unloading with botox (C, G, K, O (magnification 5) and D, H, L, P (magnification 20)). The area of magnification 20 is shown as a square in the picture above with magnification 5 i.e. B is showing the squared area in A. The black lines are marking the tendon stump.

## Discussion

In this study we investigated how two different magnitudes of daily loading affect the biomechanical and collagen properties of healing rat Achilles tendons over time. We focused on the effect on collagen orientation and its possible links to the important viscoelastic properties. The mechanical characterisation showed that the magnitude of loading is most consequential in the early healing process where creep and stiffness properties and ultimate strength (peak force) differed between the loading scenarios for up to 2 weeks post-injury. These were also the time-points where differences were seen between the treatment groups in tissue maturation measures from SAXS (Peak FWHM; D-spacing) and histological analyses. The changes in biomechanical behaviour generally diminished over time and there were not much difference between normal loading and unloading after 4 weeks of healing.

At 1 week of healing, during the first creep test (up to 5N) both creep ratio and creep magnitude were reduced in unloaded tendons while stiffness and Young’s modulus was higher compared to the loaded tendons. In contrast, during the load to failure test, stiffness and peak force was lower in the unloaded group while Young’s modulus and peak stress at failure were similar between the two groups (week 1). The decreased stiffness and peak force in the load to failure test in the unloading group is consistent with previous observations (Eliasson et al. 2009; Hammerman et al. 2018). Whereas, the higher stiffness in unloaded tendons during the 5N creep test is contradictory to other experimental findings. This could be linked to that first of all the preconditioning (10 cycles between 1-2 N) likely affected the tendons differently. The 2 N force corresponded to ∼ 20% of the maximum force of the unloaded tendons. Thus, the collagen fibrils were more strained during the preconditioning compared to in the loaded tendons where 2 N was still in the toe region leaving the fibres more crimped. Secondly, the difference in length of the tendons resulted in the 5N load being more in the linear regime in the unloaded tendons while it had yet not reached there in the loaded samples.

The large creep response at 1 week post-injury in the loaded group, together with the low stiffness properties during the 5N creep load and high stiffness at load to failure suggest that the loaded tendons contains disorganised fibres that are not fully recruited at the load levels used in the creep tests. This disorganized tissue seems to slide or align during the creep test as the creep ratio (thus strains) is very high. Over time, the loaded tendon stiffens, peak strength increases, collagen fibres mature, and creep magnitude is reduced. Tendons in the unloaded group are initially stiffer at the creep load and experience less creep. The finding that unloading displayed decreased creep properties (ratio and magnitude) is consistent with our previous study, where we observed less viscoelastic (decreased stress-relaxation and creep response) tendons as a result of reduced loading in intact tendons (Khayyeri et al. 2017).

The measured gap size between the tendon stumps were significantly larger in the loaded compared to the unloaded tendons at 1 and 2 weeks. This observation was also obvious in the histological analysis. Note that measurement of gap size became more difficult with time when the surrounding callus tissues became thicker around the stumps, especially at 4 weeks where the stumps have started to degrade as seen in the histological samples. However, it was possible by holding the sample in front of a strong light. Creep ratio was calculated based on the gap size (rather than the grip-to-grip size) because 1) we calculated that over 90% of the displacements initially occurred in the callus tissue, and 2) the gap size varied between the treatments, which would have pre-disposed the creep ratio calculations. The calliper measurements may be somewhat primitive, but the relative difference between groups still hold.

Tissue analysis, with SAXS and histology, showed that the loaded tendons had a more mature tendon tissue compared to the unloaded tendons at 1 week of healing. The loaded tendons were more mature with better collagen alignment (i.e. a lower histology score and higher D-spacing). The positive effect of loading was also observed at 2 weeks with SAXS regarding D-spacing, and collagen packing (FWHM). This discrepancy between loaded and unloaded tendons was reduced by week 4 regarding both biomechanical properties and tissue morphology. The histological analysis showed that unloaded tendons improved significantly between 1 and 4 weeks, again proposing that loading influences primarily the first weeks of tendon healing. The improved collagen orientation due to mechanical loading has been reported by other researchers (Hillin et al. 2019), but to our knowledge no other studies have used SAXS to characterize tendon healing.

The profile analysis from SAXS and histology showed that there were spatial variations in tissue maturation. Both groups had higher intrafibrillar organisation and contained more densely packed fibres (FWHM) at the sides of the callus compared to its centre. These regional differences were maintained over time, even as the collagen parameters developed. Better tissue morphology at the sides of the callus was also somewhat apparent in the unloaded tendons in the histological analysis. Additionally, the loaded tendons in the histological analysis at 1 week showed that one side of the callus had a better morphology compared to the other side. This observation could also be seen for D-spacing, intrafibrillar organisation, and packaging of fibres (FWHM) for the medial side of the callus compared to the lateral at 1 week. This suggests that the repairing tissue is not homogenous, nor heterogeneous to the degree that it appears as random, but that perhaps the repair is initiated or accelerated by processes occurring at the sides of the callus. This could be driven by local variation in mechanical stimuli across the tendon callus, or by the fact that the paratenon in Achilles tendons are thin sheaths of fibrous tissue that surround the tendon and that are also more pronounced during the repair process. Other studies have shown that the paratenon plays an important role in tendon healing, regulating growth factors (Lyras et al. 2016), proteins essential for collagen synthesis (Dyment et al. 2013), gliding resistance during motion (Momose et al. 2002) and stimulating recovery of mechanical strength (Müller et al. 2018). The observation that the healing tendon might heal differently in different region of the healing tissue is partly corroborated by other studies (e.g. Sasaki et al. 2012).

Histology showed that the loaded group contained less adipose tissue over time, which has also been found in previous studies (Hammerman et al. 2015), suggesting that loading inhibits adipocyte formation. Interestingly, the unloaded tendons at 4 weeks displayed more unorganized collagen and more adipocytes on one side of the callus, which might be related to asymmetric loading.

Although proteoglycans were found in both groups at all time points, the loaded group displayed more chondrocytes and fibrocartilaginous-type tissue, which was primarily found at the tendon stumps. This could be due to the higher compressive forces prevailing at the stump during normal gait, which promotes chondrocyte differentiation (Angele et al. 2004; Wren et al. 2000), or it could be a result of microrupture due to high strain and early signs of ossification. At week 4, the tendon stumps were less distinct suggesting that the original tissue had degraded partly into the newly formed tissue.

This study has some limitations. The magnitude of loading in the two groups has not been quantified. Botox only reduces the load-environment and new studies are needed using tools such as gait analysis and strain gauges to quantify the mechanical environment. However, it has been shown that the effect of Botox remains 4 weeks after the injection with minor reduced effect (Cichon Jr et al. 1995). Another limitation is that the SAXS measurements were performed on the entire 3D tendon rather than on a slice or a section. This gives an average two-dimensional representation over the tendon thickness and does not capture anteroposterior differences that could have been present in the tissue. However, this presents the first SAXS mapping study on healing Achilles tendon tissue, and therefore provide unique spatial data on collagen formation and distribution during healing.

In conclusion, the process of Achilles tendon healing is affected by the magnitude of loading specifically in the early weeks of healing. The strongest effects of loading were seen on collagen structure, organisation, and biomechanical properties. Loading positively affected D-spacing and the histological maturation at the first weeks post-injury. Most effects of unloading had diminished after 4 weeks of healing. Our study also found spatial variation in tissue maturation and collagen packing and organization, suggesting important different regional activity in the callus site that requires more research to unveil.

## Acknowledgements

Our beloved colleague Per Aspenberg passed away before this manuscript could be completed and has not approved this paper. He was an active contributor to this study and the experiments were finalised before he departed. The authors would like to thank the European Community’s Seventh Framework Program (FP7/2007-2013) under grant agreement no. 262348 (ESMI, European Soft Matter Infrastructure Network) that enabled beamtime at the cSAXS beamline of the Swiss Light Source at the Paul Scherrer Institut, Villigen, Switzerland. The authors are grateful for the financial support by the Marie Curie Intra-European Fellowship for Career Development (PIEF-GA-2012-626941) (HK) and the Knut and Alice Wallenberg Foundation (Wallenberg Academy Fellows 2017.0221).

## Author contributions

HK and HI conceived the project; PA designed the experiments and provided expert advice; HI was responsible for the application for synchrotron beamtime; HK, MH and PB performed the in vivo experiments, mechanical tests and histological embedding; MH and PE performed histological analysis; TN performed mechanical testing analysis; HK, MJT and MSG performed the SAXS measurements; MJT analysed the SAXS data; PE provided expert advice on data analysis and interpretation; HK, MH, MJT and HI drafted the manuscript. All authors participated in the interpretation of the results and in editing the manuscript.

## Supplementary data

**Supplementary figure 1:**
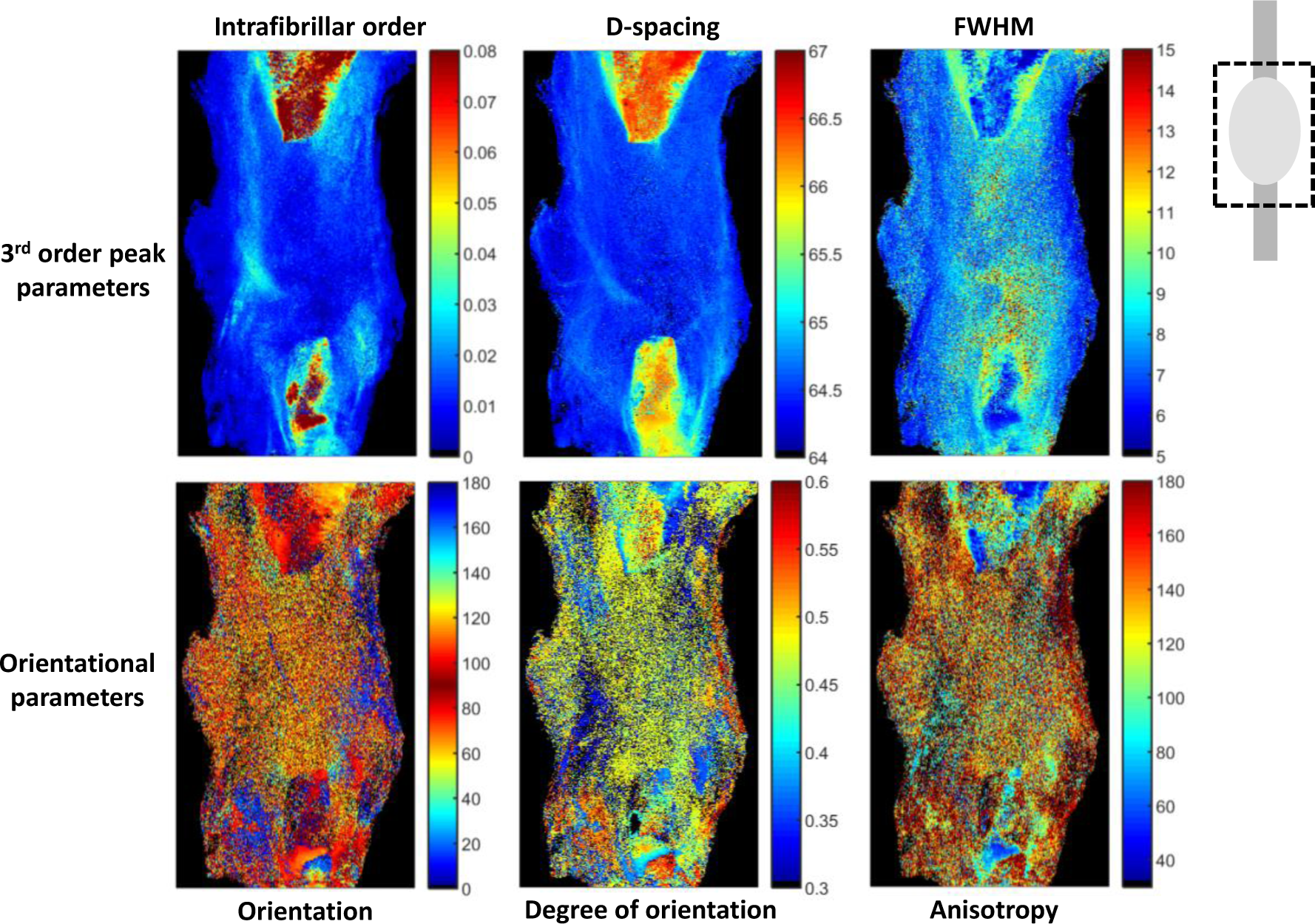
Example mapping of a healing tendon after 1 week, showing the maps of the different parameters that were quantified; D-spacing (nm), Degree of orientation (AU), anisotropy (degrees), intrafibrillar order (AU) and peak FWHM (nm).

**Supplementary figure 2:**
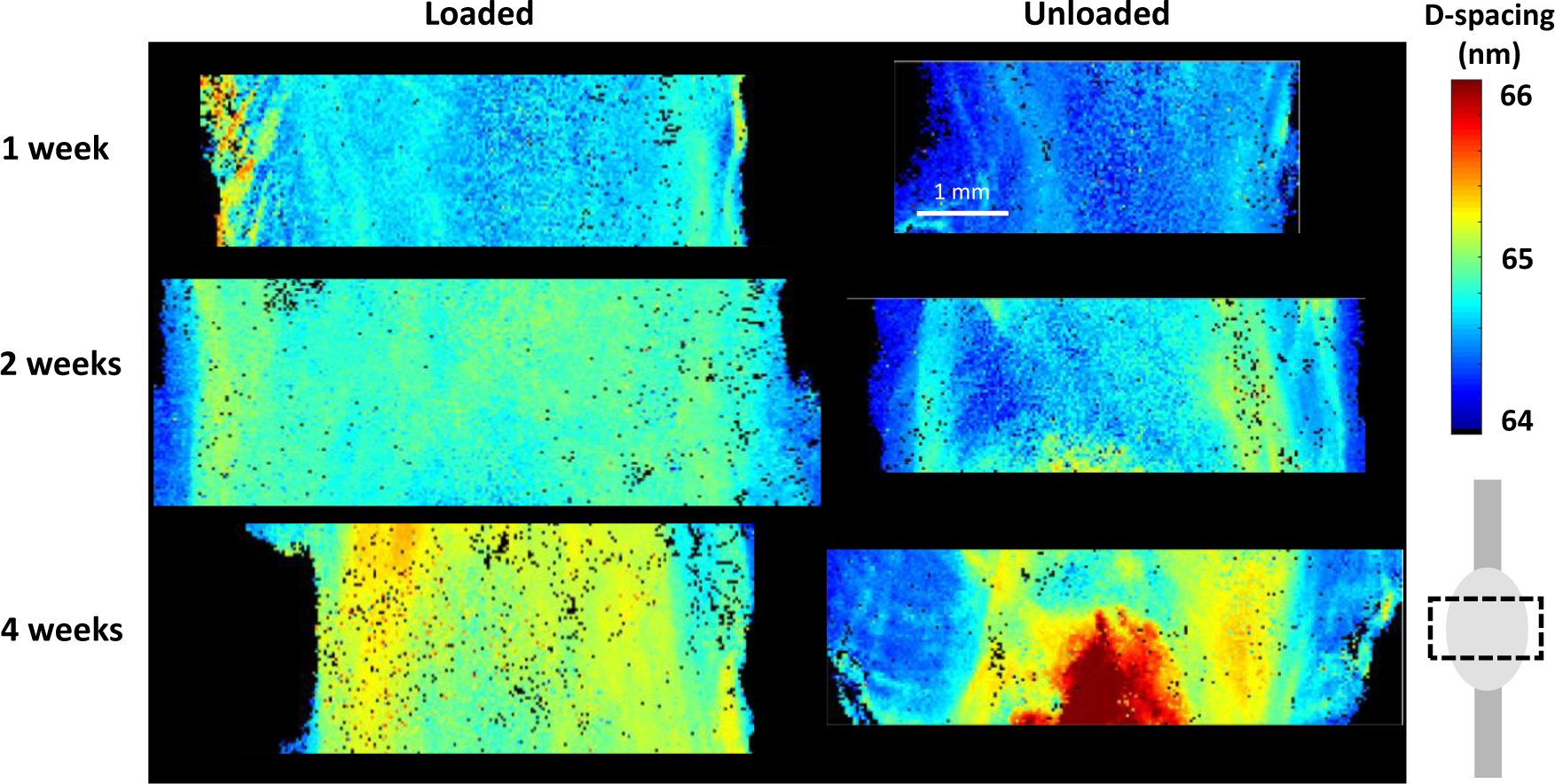
SAXS mapping of the D-spacing (nm) parameter in the callus region in representative loaded and unloaded samples at 1, 2 and 4 weeks of healing.

**Supplementary table 1:** Summary of all mechanical testing data. Data is represented as mean (standard deviation).

|  |  | W1 |  | W2 |  | W4 |  |
| --- | --- | --- | --- | --- | --- | --- | --- |
|  |  | Normal loading | Unloading | Normal loading | Unloading | Normal loading | Unloading |
|  | N | 10 | 9 | 10 | 10 | 10 | 10 |
| Callus length (mm) |  | 5.5 (1.5) | 4.1 (0.6) | 7.1 (2.7) * | 4.6 (2.3) * | 4.9 (1.5) <sup>ψ</sup> | 3.8 (1.2) |
| Cross-sectional area (mm<sup>2</sup>) |  | 23.3 (6.4) ** | 10.7 (2.9) ** | 20.8 (6.5) | 16.6 (6.1) <sup>ψ</sup> | 17.8 (3.9) | 15.6 (3.9) |
| Creep magnitude (mm) | 5N creep | 2.0 (0.3) * | 1.1 (0.1) * | 1.7 (0.5) * | 1.4 (0.4) * | 1.5 (0.3) # | 1.4 (0.4) |
|  | 12N creep | 1.7 (0.4) | - | 1.4 (0.4) | 0.6 (0.4) | 1.2 (0.2) # | 1.1 (0.3) <sup>ψ</sup> |
| Creep ratio (%) | 5N creep | 40 (16) * | 28 (3) * | 25 (6) * <sup>ψ</sup> | 33 (9) * | 33 (8) | 37(4) # |
|  | 12N | 41 (14) | - | 29 (20) | 32 (7) | 27 (6) | 29 (5) |
| Stiffness (N/mm) | 5N creep | 3.1 (0.4) ** | 4.9 (0.7) ** | 4.2 (1.4) | 4.9 (1.7) | 4.5 (2.0) | 4.4 (1.0) |
|  | 12N creep | 6.8 (1.0) | - | 10.4 (2.4) | 10.2 (2.6) | 6.1 (1.5) | 11.0 (2.8) |
|  | Load to failure | 14.5 (4.5) * | 7.8 (1.0) * | 22.5 (3.9) <sup>ψ</sup> | 21.7 (4.1) | 28.0 (4.4) ### | 25.8 (6.1) ### |
| Young's modulus (MPa) | 5N creep | 0.8 (0.4) * | 2.0 (0.7) * | 1.4 (0.5) | 1.3 (0.4) | 1.2 (0.4) | 1.1 (0.4) ## |
|  | 12N creep | 2.2 (1.1) | - | 3.1 (0.9) | 2.8 (0.9) | 2.4 (0.4) | 2.7 (0.8) |
|  | Load to failure | 3.9 (2.0) | 3.2 (1.0) | 6.9 (1.8) | 6.1 (1.7) <sup>ψψ</sup> | 7.6 (2.0) ## | 6.5 (2.2) ### |
| Peak force (N) |  | 24.1 (6.4) ** | 11.4 (1.0) ** | 46.8 (20.0) * | 27.0 (4.9) * <sup>ψ</sup> | 60.2 (17.9) ## | 55.0 (18.2) <sup>ψψ</sup> ### |
| Peak stress (MPa) |  | 1.1 (0.3) | 1.1 (0.3) | 2.2 (0.6) <sup>ψψ</sup> | 1.9 (0.6) <sup>ψ</sup> | 3.4 (0.6) <sup>ψψ</sup> ### | 3.6 (0.9) <sup>ψψψ</sup> ### |

\* $p < 0.05$ , \*\* $p < 0.01$ , \*\*\* $p < 0.001$
\* significant difference between normal loading and unloading by botox, within the same time point

**Supplementary table 2:** Summary of results from SAXS analysis from the full tendon and the separate regions of interest. Data is represented as mean (standard deviation).

|  |  | Week 1 |  | Week 2 |  | Week 4 |  |
| --- | --- | --- | --- | --- | --- | --- | --- |
|  |  | Normal loading | Unloading | Normal loading | Unloading | Normal loading | Unloading |
|  | N | 7 | 8 | 6 | 7 | 6 | 7 |
| D-spacing (nm) | Full tendon | 64.63 (0.03) | 64.54 (0.02) | 64.88 (0.02) | 64.80 (0.06) | 65.04 (0.04) | 65.09 (0.19) |
|  | Lateral | 64.57 (0.06) | 64.54 (0.05) | 64.89 (0.02) | 64.73 (0.07) | 65.05 (0.07) | 65.07 (0.19) |
|  | Centre | 64.70 (0.02) | 64.60 (0.06) | 64.86 (0.04) | 64.86 (0.06) | 65.07 (0.03) | 65.06 (0.10) |
|  | Medial | 64.70 (0.02) | 64.60 (0.06) | 64.86 (0.06) | 64.86 (0.04) | 65.07 (0.03) | 65.06 (0.10) |
| Intrafibrillar order (AU) | Full tendon | 0.017 (0.001) | 0.016 (0.002) | 0.022 (0.005) | 0.018 (0.005) | 0.048 (0.016) | 0.038 (0.020) |
|  | Lateral | 0.016 (0.002) | 0.016 (0.004) | 0.020 (0.004) | 0.015 (0.005) | 0.041 (0.014) | 0.033 (0.017) |
|  | Centre | 0.014 (0.002) | 0.012 (0.001) | 0.022 (0.004) | 0.013 (0.001) | 0.039 (0.011) | 0.037 (0.024) |
|  | Medial | 0.020 (0.003) | 0.020 (0.005) | 0.026 (0.008) | 0.023 (0.002) | 0.063 (0.027) | 0.044 (0.021) |
| Degree of orientation (AU) | Full tendon | 0.42 (0.01) | 0.44 (0.01) | 0.45 (0.02) | 0.45 (0.02) | 0.46 (0.01) | 0.46 (0.02) |
|  | Lateral | 0.42 (0.02) | 0.43 (0.02) | 0.45 (0.01) | 0.45 (0.02) | 0.46 (0.01) | 0.46 (0.02) |
|  | Centre | 0.42 (0.01) | 0.43 (0.02) | 0.44 (0.02) | 0.45 (0.02) | 0.46 (0.04) | 0.46 (0.04) |
|  | Medial | 0.43 (0.01) | 0.45 (0.02) | 0.47 (0.03) | 0.46 (0.04) | 0.45 (0.01) | 0.47 (0.02) |
| FWHM (nm) | Full tendon | 7.60 (0.23) | 7.60 (0.23) | 8.17 (0.40) | 8.86 (0.47) | 8.32 (0.25) | 8.05 (0.45) |
|  | Lateral | 7.77 (0.31) | 7.39 (0.37) | 8.17 (0.48) | 8.85 (0.36) | 8.35 (0.28) | 8.22 (0.49) |
|  | Centre | 8.03 (0.42) | 8.40 (0.35) | 8.62 (0.41) | 9.79 (0.77) | 8.60 (0.27) | 8.45 (0.44) |
|  | Medial | 6.98 (0.24) | 7.38 (0.60) | 7.69 (0.55) | 7.88 (0.48) | 8.02 (0.24) | 7.46 (0.50) |
| Anisotropy (degrees) | Full tendon | 138.2 (4.5) | 135.6 (4.9) | 142.8 (7.0) | 134.0 (3.2) | 125.0 (16.4) | 127.7 (10.6) |
|  | Lateral | 134.7 (8.8) | 133.3 (8.0) | 143.8 (2.7) | 143.2 (5.2) | 128.3 (15.0) | 131.1 (9.8) |
|  | Centre | 136.6 (6.4) | 136.3 (5.2) | 144.6 (6.9) | 140.4 (8.0) | 137.6 (17.9) | 133.9 (17.8) |
|  | Medial | 142.6 (7.8) | 137.0 (6.5) | 139.7 (15.2) | 120.1 (10.5) | 109.4 (22.4) | 117.7 (17.8) |

## References

Abrahams M (1967) Mechanical behaviour of tendonIn vitro. Medical and Biological Engineering 5:433–443

Angele P et al. (2004) Cyclic, mechanical compression enhances chondrogenesis of mesenchymal progenitor cells in tissue engineering scaffolds. Biorheology 41:335–346

Aubry S, Nueffer J-P, Tanter M, Becce F, Vidal C, Michel F (2014) Viscoelasticity in Achilles tendonopathy: quantitative assessment by using real-time shear-wave elastography. Radiology 274:821–829

Buck RC (1980) Reorientation response of cells to repeated stretch and recoil of the substratum. Experimental cell research 127:470–474

Chiquet M, Renedo AS, Huber F, Flück M (2003) How do fibroblasts translate mechanical signals into changes in extracellular matrix production? Matrix biology 22:73–80

Cichon Jr JV, Mccaffrey TV, Litchy WJ, Knops JL (1995) The effect of botulinum toxin type A injection on compound muscle action potential in an in vivo rat model. The Laryngoscope 105:144–148

Cohen R, Hooley C, McCrum N (1976) Viscoelastic creep of collagenous tissue. Journal of biomechanics 9:175–184

Dyment NA et al. (2013) The paratenon contributes to scleraxis-expressing cells during patellar tendon healing. PloS one 8:e59944

Eliasson P, Andersson T, Aspenberg P (2009) Rat Achilles tendon healing: mechanical loading and gene expression. Journal of applied physiology 107:399–407

Eliasson P, Fahlgren A, Pasternak B, Aspenberg P (2007) Unloaded rat Achilles tendons continue to grow, but lose viscoelasticity. Journal of applied physiology 103:459–463

Freedman BR, Rodriguez AB, Hillin CD, Weiss SN, Han B, Han L, Soslowsky LJ (2018a) Tendon healing affects the multiscale mechanical, structural and compositional response of tendon to quasistatic tensile loading. Journal of The Royal Society Interface 15:20170880

Freedman BR et al. (2018b) Dynamic Loading and Tendon Healing Affect Multiscale Tendon Properties and ECM Stress Transmission. Scientific reports 8:10854

Hammerman M, Blomgran P, Ramstedt S, Aspenberg P (2015) COX-2 inhibition impairs mechanical stimulation of early tendon healing in rats by reducing the response to microdamage. Journal of applied physiology 119:534–540

Hammerman M, Dietrich-Zagonel F, Blomgran P, Eliasson P, Aspenberg P (2018) Different mechanisms activated by mild versus strong loading in rat Achilles tendon healing. PloS one 13:e0201211

Hillin CD, Fryhofer GW, Freedman BR, Choi DS, Weiss SN, Huegel J, Soslowsky LJ (2019) Effects of Immobilization Angle on Tendon Healing after Achilles Rupture in a Rat Model. Journal of Orthopaedic Research®

Khayyeri H et al. (2017) Achilles tendon compositional and structural properties are altered after unloading by botox. Scientific reports 7:13067

Killian ML, Cavinatto L, Galatz LM, Thomopoulos S (2012) The role of mechanobiology in tendon healing. Journal of shoulder and elbow surgery 21:228–237

Kukreti U, Belkoff SM (2000) Collagen fibril D-period may change as a function of strain and location in ligament. Journal of biomechanics 33:1569–1574

Lavagnino M, Wall ME, Little D, Banes AJ, Guilak F, Arnoczky SP (2015) Tendon mechanobiology: current knowledge and future research opportunities. Journal of Orthopaedic Research 33:813–822

Lyras DN, Kazakos K, Tilkeridis K, Kokka A, Ververidis A, Botaitis S, Agrogiannis G (2016) Temporal and spatial expression of tgf-b1 in the early phase of patellar tendon healing after application of platelet rich plasma. Archives of Bone and Joint Surgery 4:156

Momose T, Amadio PC, Zobitz ME, Zhao C, An KN (2002) Effect of paratenon and repetitive motion on the gliding resistance of tendon of extrasynovial origin. Clinical Anatomy: The Official Journal of the American Association of Clinical Anatomists and the British Association of Clinical Anatomists 15:199–205

Müller SA, Evans CH, Heisterbach PE, Majewski M (2018) The role of the paratenon in Achilles tendon healing: a study in rats. The American journal of sports medicine 46:1214–1219

Rigby BJ, Hirai N, Spikes JD, Eyring H (1959) The mechanical properties of rat tail tendon. The Journal of general physiology 43:265–283

Turunen MJ, Khayyeri H, Guizar-Sicairos M, Isaksson H (2017) Effects of tissue fixation and dehydration on tendon collagen nanostructure. Journal of structural biology 199:209–215

Wall ME et al. (2016) Cell signaling in tenocytes: response to load and ligands in health and disease. In: Metabolic Influences on Risk for Tendon Disorders. Springer, pp 79–95

Wren TA, Beaupre GS, Carter DR (2000) Mechanobiology of tendon adaptation to compressive loading through fibrocartilaginous metaplasia. Journal of rehabilitation research and development 37:135–144

